## Extended Data Tables and Figures for "Contributions and synaptic basis of diverse cortical neuron responses to task performance"

| A. Model Summary | |
|---|---|
| **Populations** | Spiking excitatory input units, excitatory output units, and inhibitory units. One analog readout node receives inputs from output units. |
| **Connectivity** | Random sparse connectivity (fixed probability) |
| **Neuron Model** | Leaky integrate-and-fire |
| **Synapse Model** | Current-based exponential synapses. |
| **Plasticity** | 1) Excitatory-to-excitatory and inhibitory-to-excitatory STDP.<br>2) FORCE modifications of the connections to the readout node.<br>3) Bias current to set the inhibitory firing rate. |

| B. Populations | | |
|---|---|---|
| **Variable** | **Description** | **Value** |
| $N_E$ | Number of excitatory units | 800 |
| $N_{\text{in}}$ | Subpopulation of excitatory units that receive external inputs | 200 |
| $N_{\text{out}}$ | Subpopulation of excitatory units that project connections to the readout node | 600 |
| $N_I$ | Number of inhibitory units | 200 |

| C. Neuron Model | | |
|---|---|---|
| **Variable** | **Description** | **Value** |
| $V_r$ | Resting membrane potential | -65 mV |
| $V_{th}$ | Threshold membrane potential | -55 mV |
| $\tau$ | Membrane time constant | 20 ms |
| $\tau_E$ | EPSP time constant | 20 ms |
| $\tau_I$ | IPSP time constant | 20 ms |
| $R_I$ | Inhibitory baseline firing rate target | 20 spikes/s |
| $\eta_r$ | Bias current learning rate | 0.005 |
| $I_{\text{in}}$ | Stimulus input current | 2.0 pA |

| D. STDP | | |
|---|---|---|
| **Variable** | **Description** | **Value** |
| $p_{\text{con}}$ | Synaptic connection probability | 5% |
| $W_{0\,E}$ | Initial excitatory synaptic outputs | 1.4 |
| $W_{0\,I}$ | Initial inhibitory synaptic outputs | 1.4 |
| $\tau_+$ | Excitatory-to-excitatory LTP timescale | 20 ms |
| $\tau_-$ | Excitatory-to-excitatory LTD timescale | 20 ms |
| $A$ | Excitatory-to-excitatory LTP magnitude | 1e-3 |
| $B$ | Excitatory-to-excitatory LTD magnitude | 1.05e-3 |
| $\beta_E$ | Excitatory-to-excitatory heterosynaptic balancing parameter | 5.62e-4 |
| $\delta_E$ | Excitatory-to-excitatory heterosynaptic strengthening parameter | 1e-4 |
| $\tau_I$ | Inhibitory-to-excitatory STDP timescale | 5 ms |
| $R_E$ | Inhibitory-to-excitatory target excitatory rate | 10 spikes/s |
| $\alpha$ | Inhibitory-to-excitatory STDP LTD magnitude | 0.1 |
| $\eta_I$ | Inhibitory-to-excitatory STDP overall magnitude | 1e-3 |
| $\beta_I$ | Excitatory-to-excitatory heterosynaptic balancing parameter | 5.62e-4 |
| $\delta_I$ | Excitatory-to-excitatory heterosynaptic strengthening parameter | 1e-4 |

| E. FORCE | | |
|---|---|---|
| **Variable** | **Description** | **Value** |
| $\tau_{\text{out}}$ | Readout node time constant | 100 ms |
| $T_{\text{FORCE}}$ | Average time between FORCE output weight updates | 4 ms |
| $Q$ | Feedback strength | 2.0 |

**Extended Data Table 1: RNN parameters.** Default RNN parameters. All networks used these parameters unless otherwise stated.

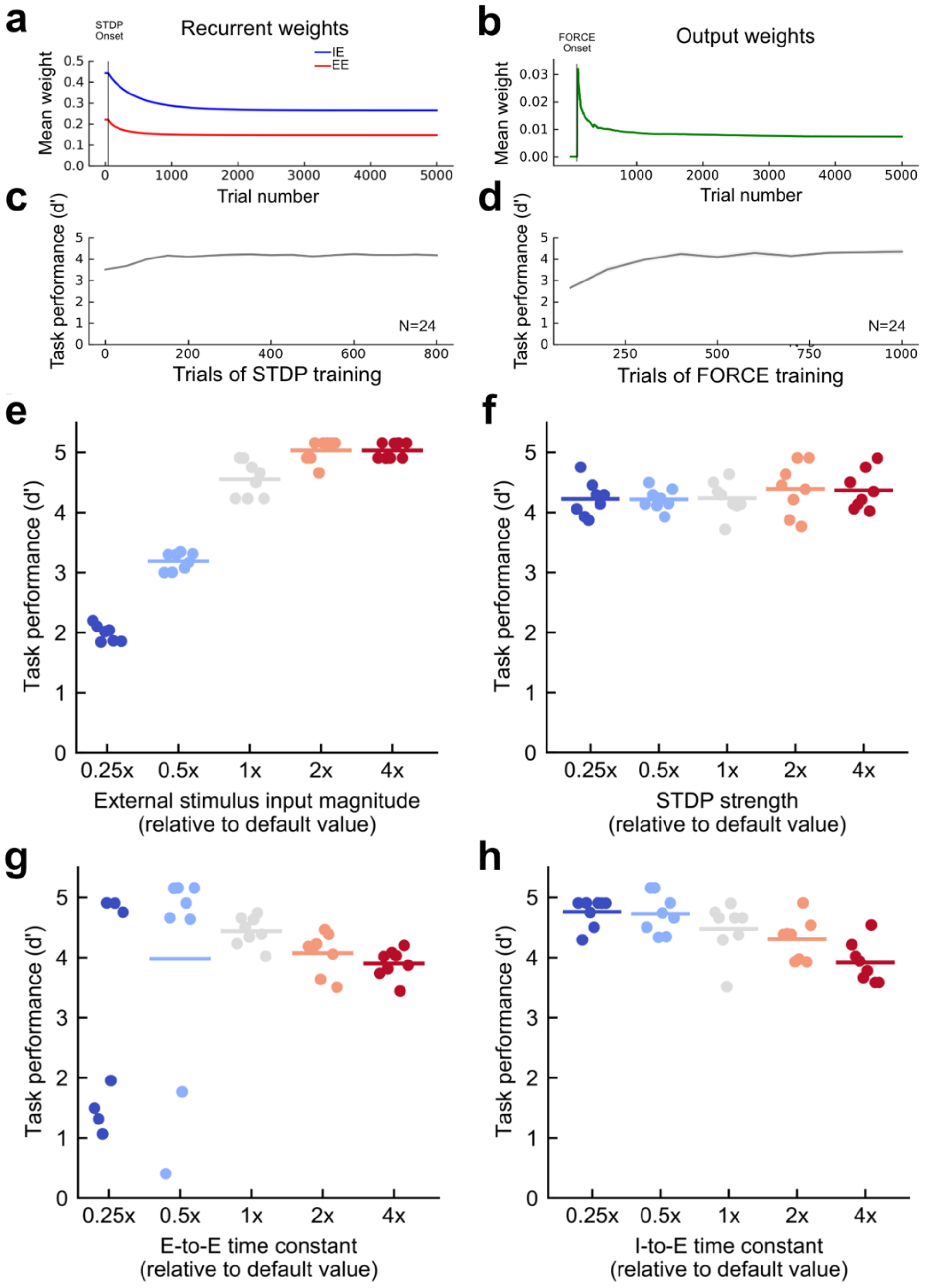


**Extended Data Figure 1. Network training and parameter selection. a**, Time course of recurrent network weights modified by STDP over training **b,** Time course of output weights modified by FORCE over training. **c,** Asymptotic task performance (d') vs. number of STDP trials during training. **d,** Asymptotic task performance (d') vs. number of FORCE trials during training. In these networks, STDP and FORCE plasticity mechanisms were active sequentially. In **c** and **d**, trials where STDP and FORCE plasticity mechanisms were not active simultaneously with STDP occurring first. **e,** Asymptotic task performance (d') vs. strength of external currents injected into excitatory input units. Circles, task performance of individual networks (N=8 per group); horizontal lines, means. Default value was chosen because it allowed for high performance while permitting some errors. **f,** Asymptotic task performance (d') vs. STDP strength. **g** and **h,** Asymptotic behavioral performance (d') for excitatory-to-excitatory and inhibitory-to-excitatory time constants relative to network defaults.

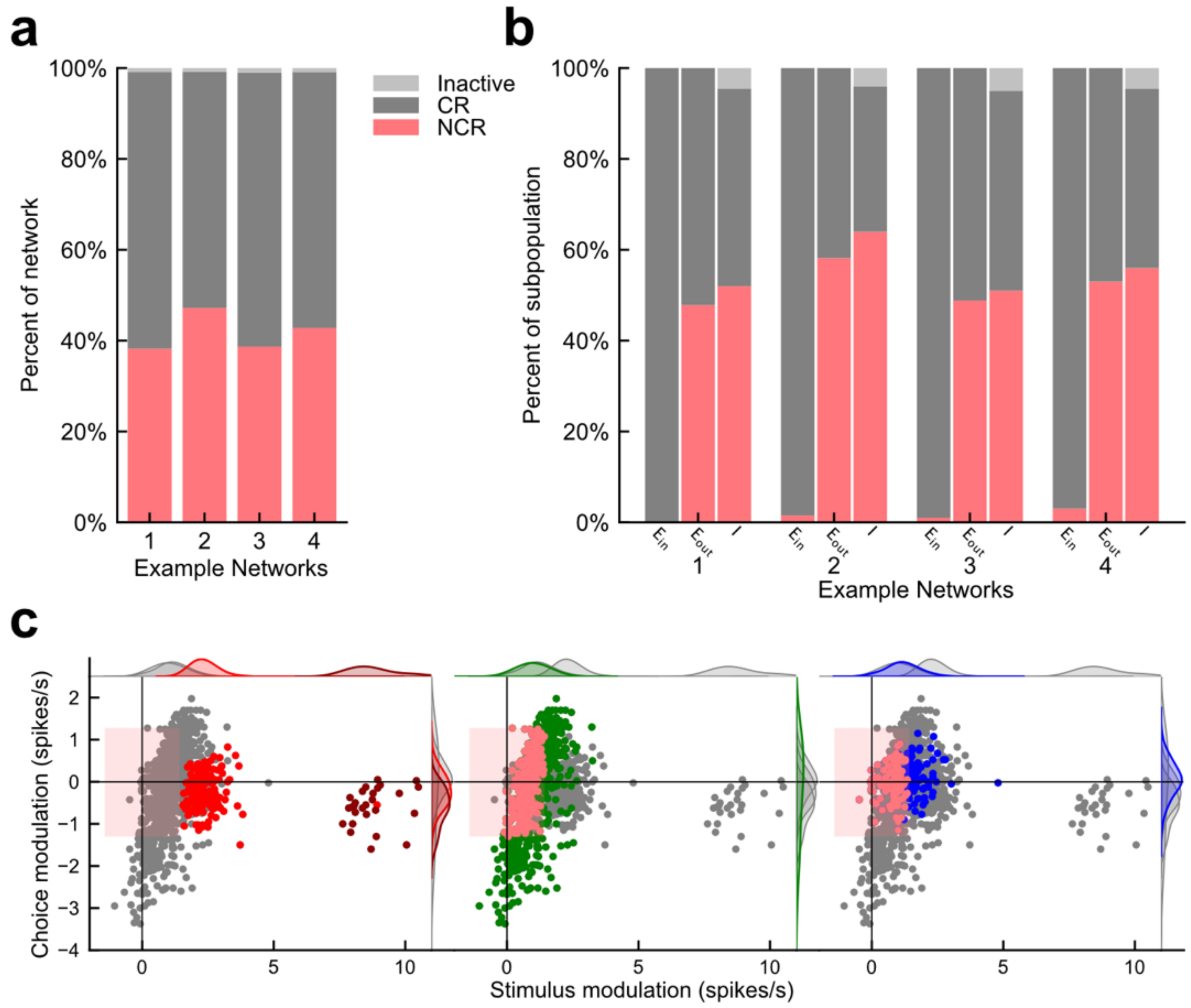


**Extended Data Figure 2. Example population statistics for RNN model. a,** Example population statistics for four example networks. Non-classically responsive units (NCR, red), classically responsive units (CR, dark grey), inactive units (inactive light grey). **b,** Population statistics for 4 networks broken out by subpopulation: excitatory input units ($E_{in}$), excitatory output units ($E_{out}$), Inhibitory units ($I$). **c,** Choice versus stimulus modulation for each unit in an example post-STDP network. All units are shown in each panel with different subpopulations highlighted in each. Left, excitatory input units are highlighted, non-target selective units (red), target selective units (maroon). Middle, output units highlighted in green. Right, inhibitory units highlighted in blue. Non-classically responsive units are shown in light red on each panel. Light red rectangle overlay corresponds to the statistical criteria to designate non-classically responsive units. Left, excitatory input units are highlighted, non-target selective units (red), maroon (target) selective units. As expected these units are highly stimulus modulated, but less modulated during the choice period. Middle, output units highlighted in green. These units span a wide range of both stimulus and choice modulation values. Right, inhibitory units highlighted in blue.

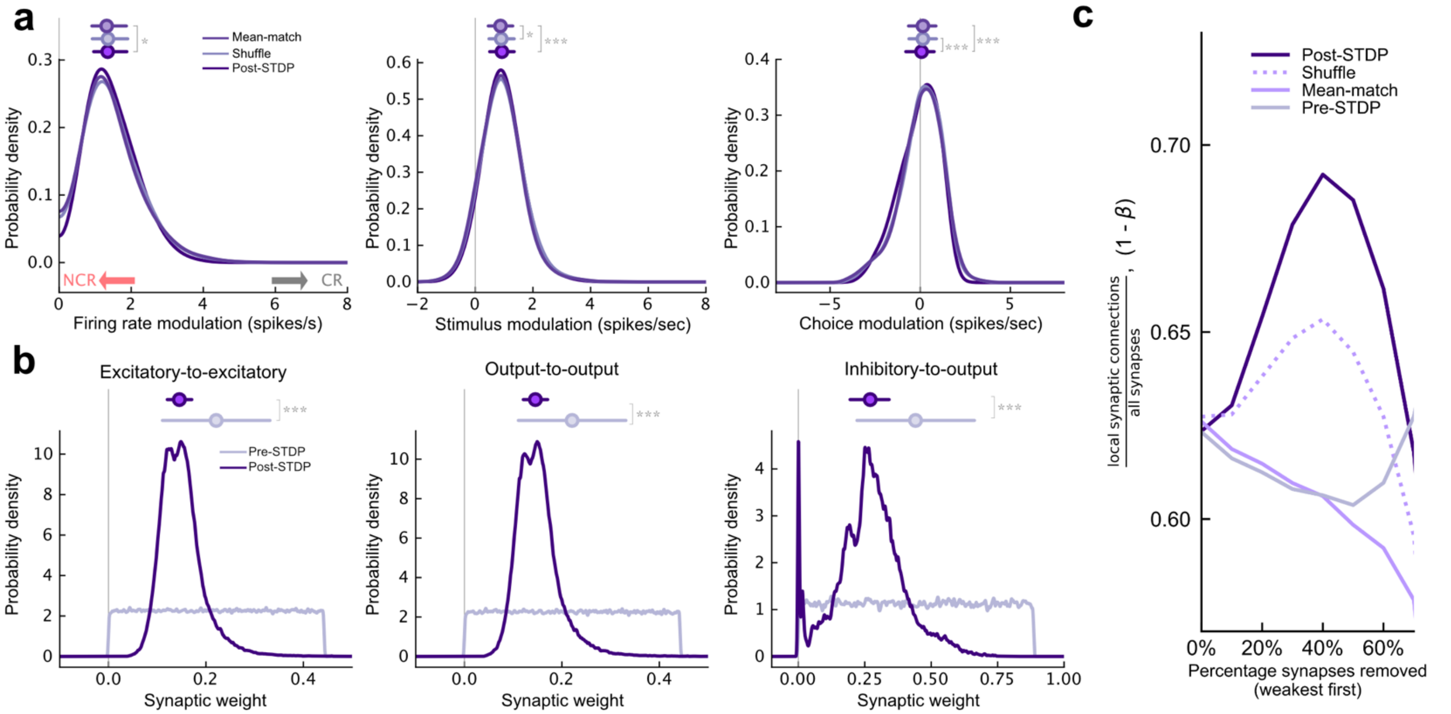


**Extended Data Figure 3. Effect of STDP on network architecture and response profiles**. **a,** Overall firing rate modulation (left), stimulus period firing rate modulation (center), choice period firing rate modulation (right) distributions for post-STDP networks versus mean-matched and shuffled controls with inactive units excluded. **b,** Distribution of synaptic weights pre-STDP (light purple) and post-STDP (purple). Summary statistics above distributions represent median and interquartile range. Left, distributions for all excitatory-to-excitatory connections. Center, excitatory output-to-output connections only. Right, inhibitory-to-excitatory connections. **c,** Fraction of local synaptic connections for Pre-STDP and Post-STDP networks as well as shuffle and mean-match controls. This number was calculated by comparing the mean shortest path length between all units in the network and comparing this to the value generated by small-world networks with various fractions of random, non-local connections (β). Because the small-world criteria does not account for synaptic strength different percentages of the weakest synapses were removed. e.g., when 40% of the weakest synapses were removed Pre-STDP networks were comparable to networks where ~60% of the synaptic connections were local whereas Post-STDP networks were comparable to networks where ~70% were local. This increase is consistent with STDP inducing a 'small-world' structure in the recurrent weights.

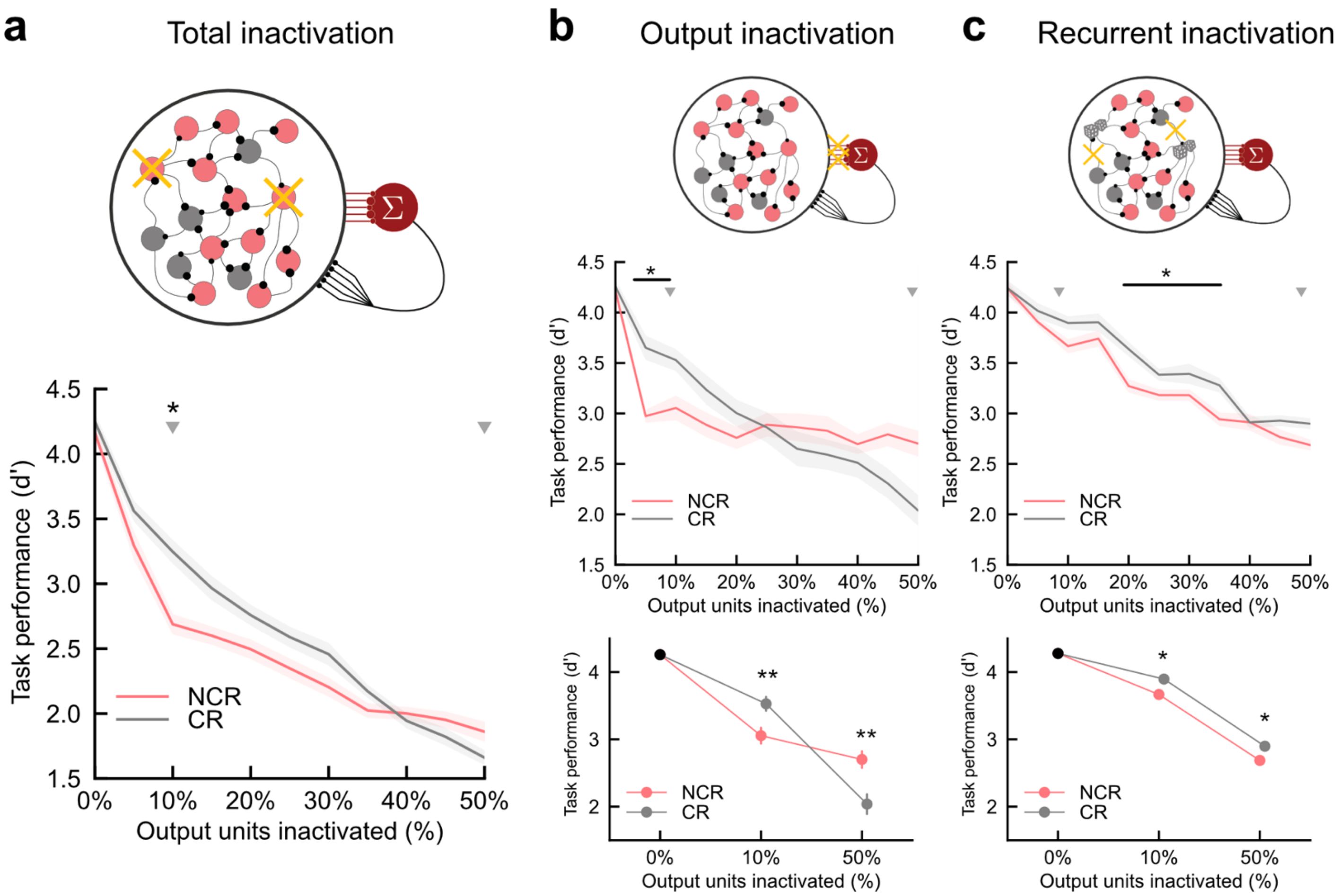


**Extended Data Figure 4. Classically and non-classically responsive units contribute to task performance via distinct mechanisms. a,** Task performance as a function of output units inactivated for non-classically responsive units only (red, NCR) and classically responsive units only (grey, CR). Lines and shading represent mean and SEM, respectively. Grey triangles at 10% (n=60) and 50% (n=300) indicate values shown in Fig 3d. **b,** Task performance as a function of output weight inactivation only (leaving recurrent connections intact). Removing output connections from 60 units i.e. 10% most non-classically responsive Δd' = -1.24 vs. 10% most classically responsive Δd' = -0.57, $p < 10^{-5}$, Mann-Whitney U two-sided U test with Bonferroni correction **c**, Same as **b**, but for recurrent perturbation only leaving output connections intact. Disabling recurrent connections of 60 units i.e. 10% most non-classically responsive output units Δd' = -0.74 vs. 10% most classically responsive units Δd' = -0.43,  p=0.016, Mann-Whitney U two-sided with Bonferroni-correction.

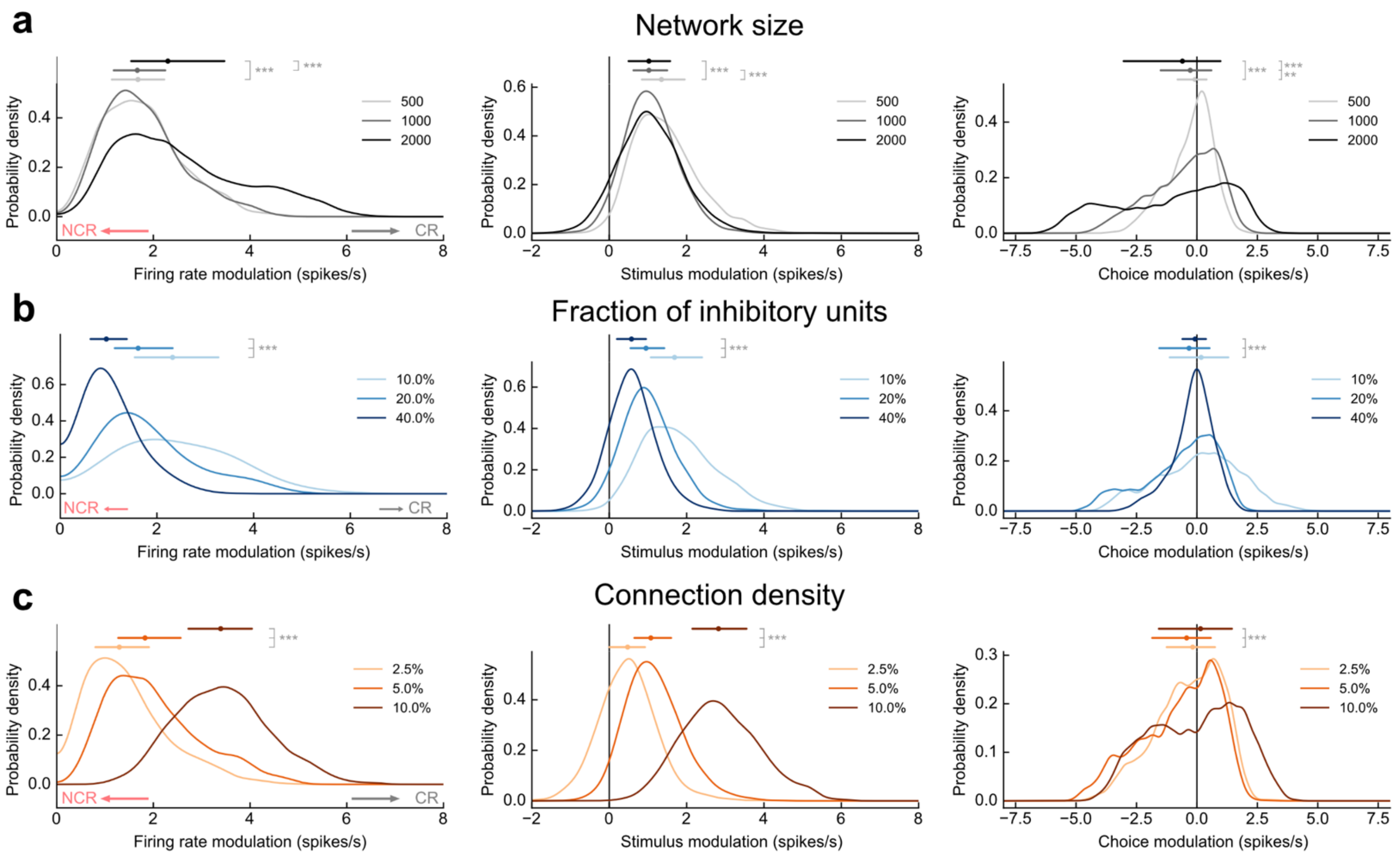


**Extended Data Figure 5. Effect of network parameters on response profile distributions. a,** Probability density of firing rate modulation for individual output units in networks of varying size (n = 500, 1000, 2000) with inhibitory fraction (f = 20%) and connection probability ($p_{con}$ = 5%) held constant. Small values of the firing rate modulation correspond to non-classical response profiles; high values correspond to classical response profiles. Summary circles and bars above distributions represent median and interquartile range, respectively. Left, firing rate modulation, $p < 10^{-5}$ for 2000 units vs 1000 and 500, Mann-Whitney two-sided U test with Bonferroni correction. Middle, stimulus modulation only, p=0.0016 for 500 vs 1000, $p<10^{-5}$ for 2000 vs. 500 and 1000. Right, Choice modulation only, $p < 10^{-5}$ all comparisons, Levene's test with Bonferroni-correction. **b,** Same as **a** except for networks of varying inhibitory fraction (f = 10%, 20%, 40%) with total network size (n = 1000) and connection probability ($p_{con}$ = 5%) held constant. Left, firing rate modulation $p < 10^{-5}$ all comparisons, Mann-Whitney two-sided U test with Bonferroni correction**.** Middle, stimulus modulation, $p < 10^{-5}$ all comparisons, Mann-Whitney two-sided U test with Bonferroni correction. Right, choice modulation, $p < 10^{-5}$ all comparisons, Levene's test with Bonferroni-correction. **c,** Same as **a** except with networks of varying connection probability ($p_{con}$ = 2.5%, 5%, 10%) with total network size (n = 1000) and inhibitory fraction held constant (f = 20%). Left, firing rate modulation, $p < 10^{-5}$ all comparisons, Mann-Whitney two-sided U test with Bonferroni correction. Middle, stimulus modulation, $p < 10^{-5}$ all comparisons**,** Mann-Whitney two-sided U test with Bonferroni correction. Right, choice modulation, $p < 10^{-5}$ all comparisons**,** Mann-Whitney two-sided U test with Bonferroni correction.

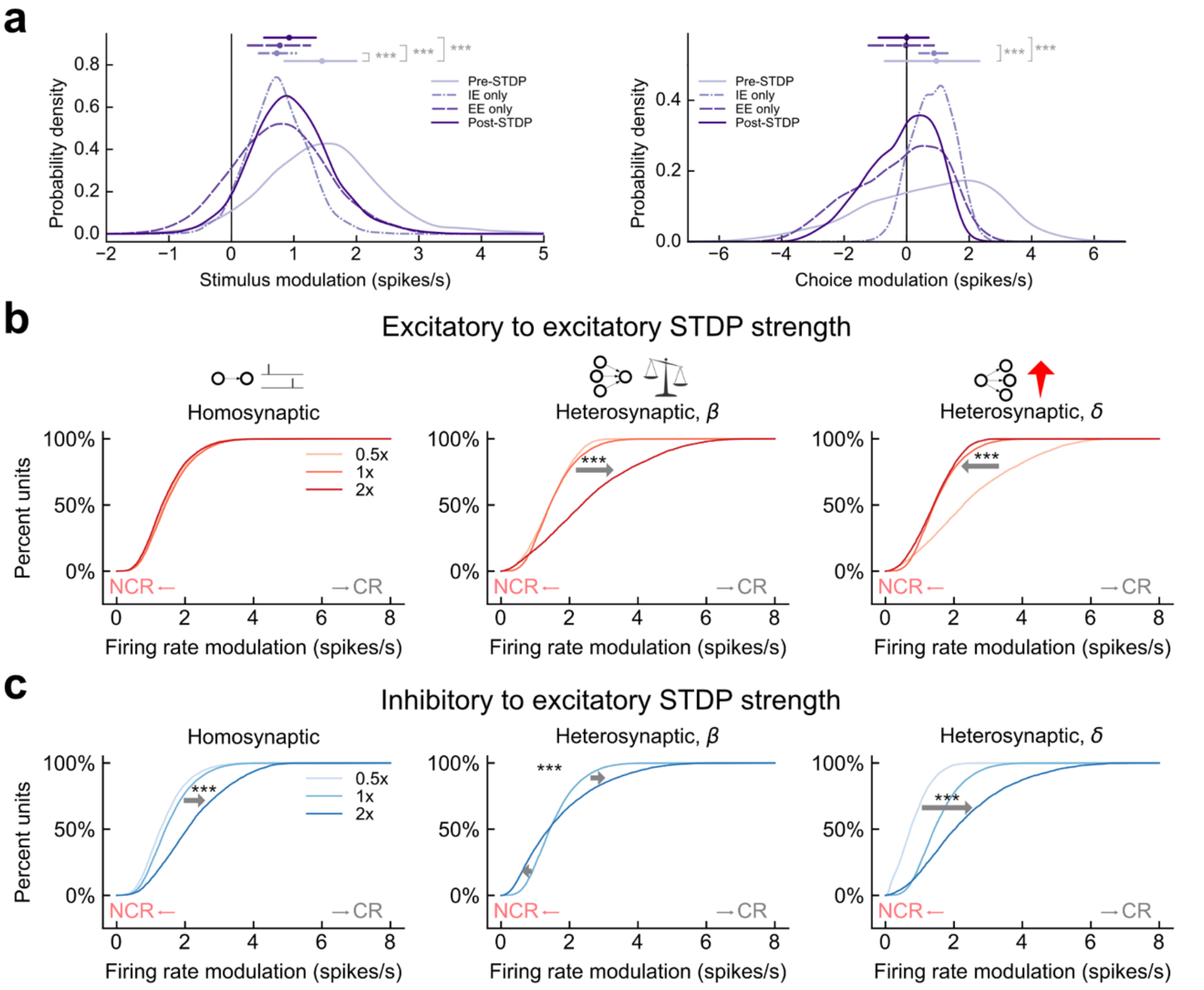


**Extended Data Figure 6. Effect of specific STDP mechanisms on response profile distributions. a,** Same as experiment **Fig. 3e** with stimulus and choice related firing rate modulation distributions displayed separately. Left, stimulus median shift relative to pre-STDP, post-STDP: Δstimulus = -0.52 spikes/s**,** IE only: Δstimulus = -0.70 spikes/s, EE only: Δstimulus = -0.67 spikes/s, $p < 10^{-5}$ for all comparisons to pre-STDP, Mann-Whitney U test two-sided Bonferroni-correction. Right, choice median shift relative to pre-STDP, post-STDP: Δchoice = -0.95 spikes/s, $p < 10^{-5}$, IE only: Δchoice = -0.07 spikes/s, p = 0.078, EE only: Δchoice = -0.97 spikes/s, $p<10^{-5}$, Mann-Whitney U test two-sided Bonferroni-correction. **b,** Cumulative probability distribution of firing rate modulation for individual output units when excitatory-to-excitatory homosynaptic (left), heterosynaptic balancing (center), and heterosynaptic enhancement (right) mechanism strengths varied relative to network defaults. $p < 10^{-5}$ for comparisons 2x to 0.5x strength, Kolmogorov-Smirnov test, Bonferroni-correction. **c,** Same as **b** but for inhibitory-to-excitatory plasticity mechanisms. $p < 10^{-5}$ for all comparisons 2x to 0.5x strength, Kolmogorov-Smirnov test, Bonferroni-correction.

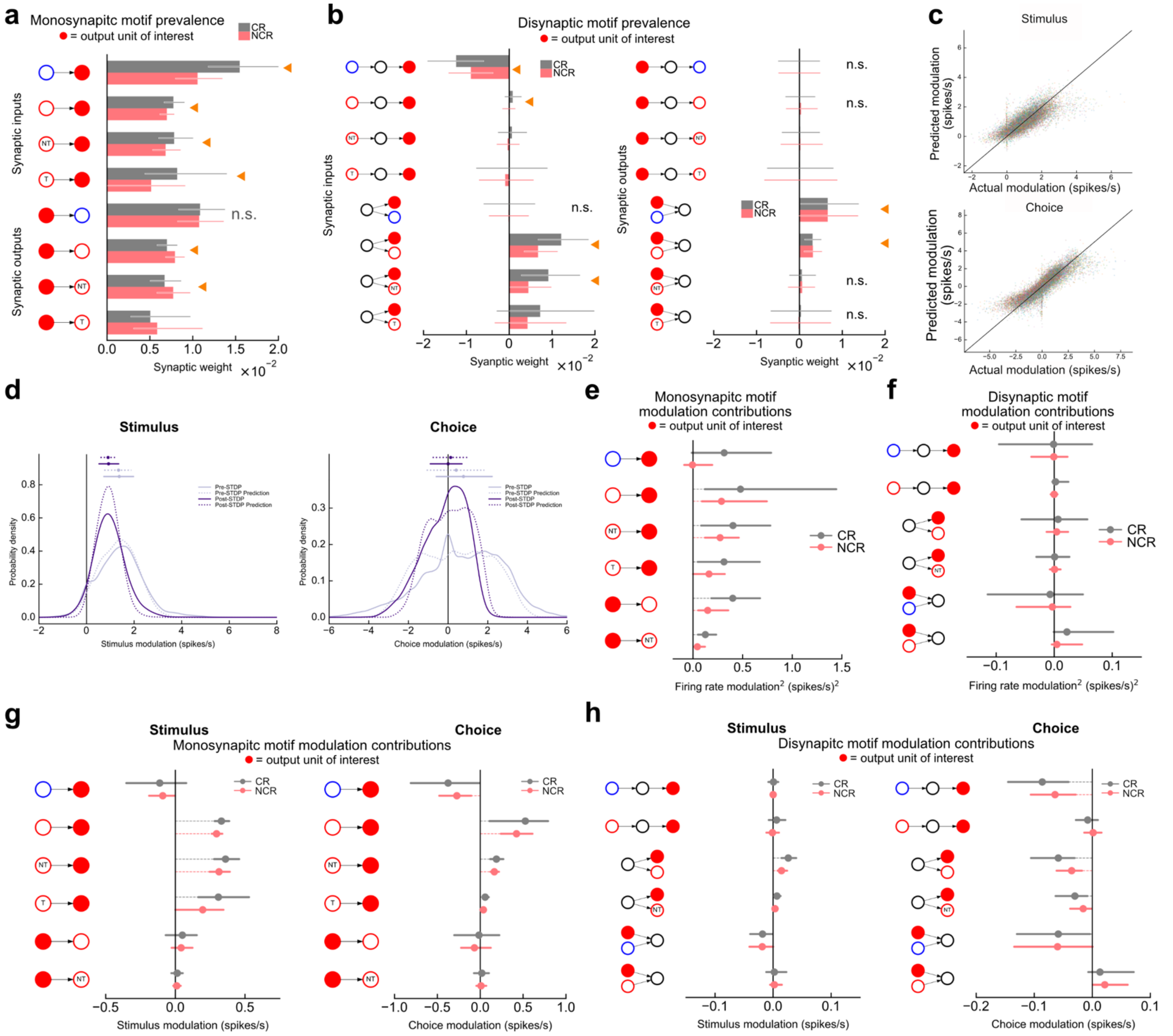


**Extended Data Figure 7. Validation of statistical motif model. a,** Prevalence of all monosynaptic motifs between individual output units and all subpopulations for non-classically responsive units (NCR, red) and classically responsive units (CR, grey). Bars and lines represent medians and interquartile range, respectively. **b**, Same as **a** but for all disynaptic motifs between individual output units and all subpopulations. **c**, 30-fold cross-validated predictions for stimulus-related firing rate modulation (top) and choice-related firing rate modulation (bottom) using statistical motif model versus actual firing rate modulation for n = 14,400 output units across 24 networks (pre-STDP, excitatory-to-excitatory STDP only, inhibitory-to-excitatory STDP only, and post-STDP). Randomly assigned colors represent different test folds. **d**, Probability densities of individual output unit stimulus-related (left) and choice-related (right) firing rate modulation for pre-STDP (light purple) and post-STDP (purple) networks and predictions (dotted lines) based on statistical motif model. Summary circles and bars above represent median and interquartile range, respectively. **e**, Contributions of individual monosynaptic motifs to single-unit firing rate modulation$^2$ for non-classically responsive units (NCR, red) and classically responsive units

(CR, grey). Circles and bars represent median and interquartile range, respectively. **f**, Same as **e** but for disynaptic motifs. **g,** Contributions of individual monosynaptic motifs to single-unit stimulus-related firing rate modulation (left) and choice-related firing rate modulation (right) for non-classically responsive units (NCR, red) and classically responsive units (CR, grey). Circles and bars represent median and interquartile range, respectively. **h**, Same as **g** but for disynaptic motifs.

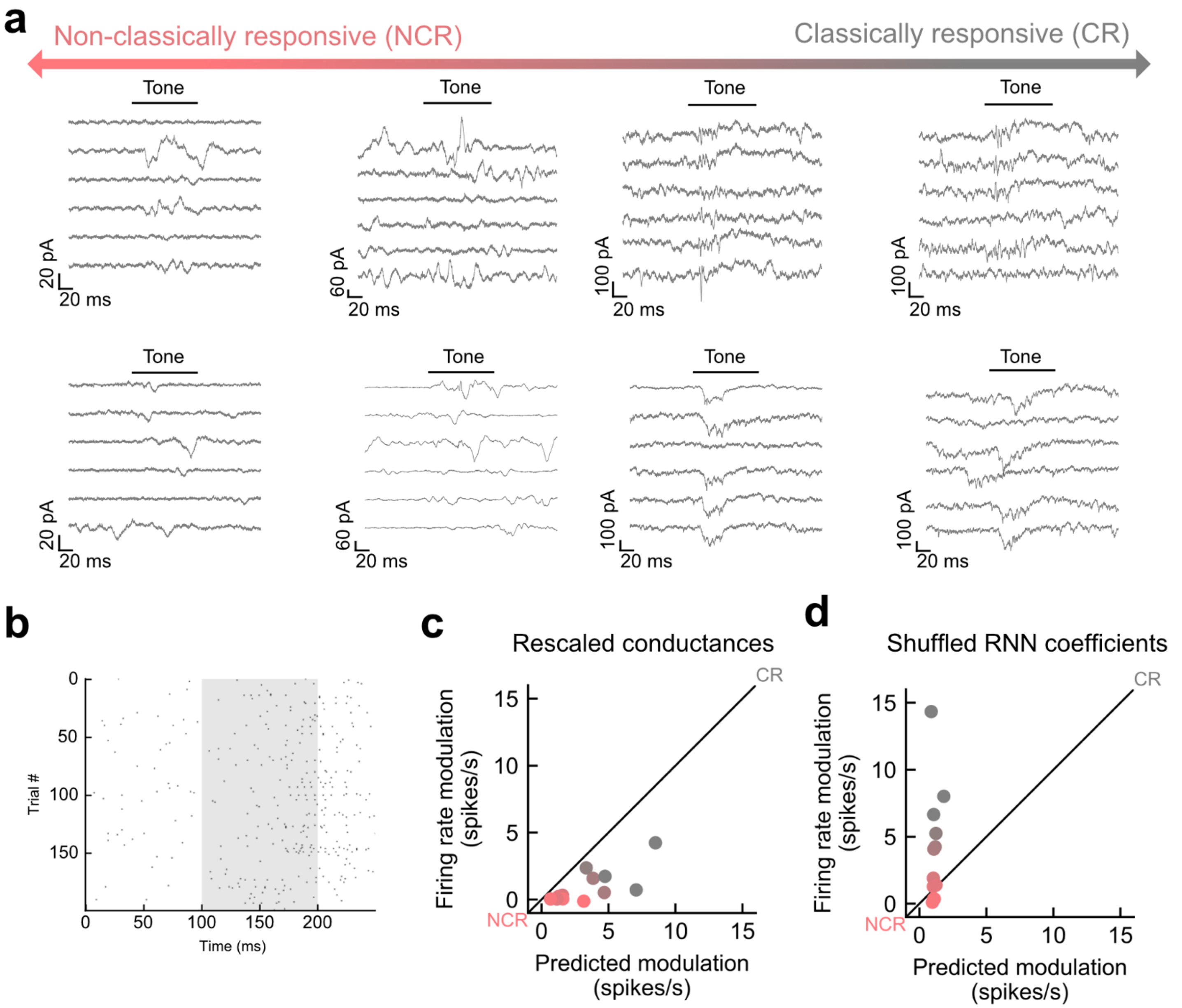


**Extended Data Figure 8. Validation of RNN-derived modulation predictions. a,** Example recordings from 4 cells spanning non-classically responsive (left) to classically responsive neuron (right). **b,** Example spiking raster from a single neuron generated using a leaky integrate-and-fire model. Trial-by-trial excitatory and inhibitory conductance dynamics were sampled randomly from conductance dynamics recorded using whole-cell voltage-clamp. **c,** Control experiment in which average conductance value used for prediction in **Fig. 5h** were left undisturbed but trial-by-trial conductance dynamics were rescaled when simulating output spikes. Under these conditions, the RNN-derived predictions deviate from simulation results indicating that the RNN-derived coefficients capture non-trivial features of the trial-by-trial dynamics *in vivo*. **d,** Control experiment in which predictions were based on RNN data that randomly shuffled the modulation values of RNN units to destroy any possible relationship between local synaptic structure and modulation. These 'shuffled RNN coefficients' were unable to predict the firing rate modulation of neurons *in vivo*.
